# Machine Learning Models of Breast Cancer Risk Prediction

**DOI:** 10.1101/723304

**Authors:** Md. Mohaimenul Islam, Tahmina Narin Poly

## Abstract

Breast cancer is the most common cancer in women both in the developed and less developed world. Early detection based on clinical features can greatly increase the chances for successful treatment. Our goal was to construct a breast cancer prediction model based on machine learning algorithms. A total of 10 potential clinical features like age, BMI, glucose, insulin, HOMA, leptin, adiponectin, resistin, and MCP-1 were collected from 116 patients. In this report, most commonly used machine learning model such as decision tree (DT), random forest (RF), K-nearest neighbors (KNN), support vector machine (SVM), logistic regression (LR), and artificial neural network (ANN) models were tested for breast cancer prediction. A repeated 10-fold cross-validation model was used to rank variables on the randomly split dataset. The accuracy of DT, RF, SVM, LR, ANN, and KNN was 0.71, 0.71, 0.77, 0.80, 0.81, and 0.86 respectively. However, The KNN model showed most higher accuracy with area under receiver operating curve, sensitivity, and specificity of 0.95, 0.80, 0.91. Therefore, identification of breast cancer patients correctly would create care opportunities such as monitoring and adopting intervention plans may benefit the quality of care in long-term.

## Introduction

Breast cancer (BC) is the most common cancer in women, and major health concern both in the developed and less developed world [1]. To detect BC at an early stage, the cure is possible but undetected at the early stage can lead to a fatal stage, even death. A major challenge is that clinical BC diagnosis is based solely on screening mammography which requires resources that are not fully available in low-and-middle-income countries [2]. Lots of efforts have already been taken to detect BC at an early stage but they did not work well. several risk factors often influence the development of BC; although, most of these do not directly cause BC. Knowing these risk factors may help patients make more informed lifestyle and health care choices. Timely recognition and treatment patients with BC would help to reduce the risk of mortality and complications and slow down disease progression. Taken together these issuers highlight a major need for early diagnosis and intervention to alleviate the impact of BC [3].

A barrier to timely recognition and management of BC is the long clinically silent phase of the disease. Patients with BC tend to be asymptomatic in the early stage, resulting in generally low awareness of the disease [4]. A woman with only a high school education has much lower screening rates than those with a college education. Additionally, a woman without insurance coverage also is screened much less frequently than those with insurance. To improve awareness at both patients and clinicians level is essential to improve the quality of care. It is therefore indispensable to get screening each year and do some diagnostic testing including serum insulin, leptin, HOMA, adiponectin, resistin, MCP-1. Crisostomo J et al. [5] reported that these variables are responsible for obesity-associated breast cancer. Furthermore, Santillan-Benitez JG, et al. [6] were assessed this variable as a biomarker for breast cancer.

Machine learning based prediction of clinical outcomes can be used for appropriate decision making and can lead to better patient care. Machine learning prediction model can accurately predict which women should undergo biopsy and help to reduce missing a breast cancer patient. ML also a great advantage over traditional statistical models including high power and accuracy to predict disease. To our knowledge, there is no specific algorithm that performs better for the prediction model. We, therefore, conducted most commonly used algorithms and compared their performance in the prediction of breast cancer.

## Methods

### Data collection

Data were collected from the UCI machine learning repository database. Protected personal health information was removed for the purpose of this research. Because it analyzed de-identified data, this study was excepted from ethics review by the Ethical Committee of CHUC and all participants gave their written informed consent prior to entering the study [7].

Patients information for this study was extracted from women newly diagnosed with breast cancer (BC) at the Gynecology Department of the University Hospital Center of Coimbra (CHUC) between 2009 and 2013. All patients were diagnosed breast cancer by standard protocol. Additionally, healthy patients were eligible to enroll in this study if they had had no prior cancer treatment and were free from any infection or other acute diseases or comorbidities at the time of enrolment. Therefore, patients with treatment before the consultation were excluded. Additionally, 38 participants were excluded due to having BMI above 40 kg/m2 or due to the absence of at least one of the quantitative variables. The actual goal of this study was then to assess hyper-resistinemia and metabolic dysregulation in breast cancer. Finally, a total of 64 women with BC and 52 healthy volunteers was included in the present study.

### Outcome Definition and identification technique

All the breast cancer patients were checked and assessed by a positive mammography and histological test. However, the tissue of breast cancer patients was collected by mastectomy or tumorectomy. Trained pathologists at the anatomic pathology department were responsible to evaluate and identify BC patient’s tumor type, grade and size and lymph node. Additionally, the TNM classification of the malignant tumors was used to confirm cancer stage. Also, estrogen and progesterone receptors status, and HER-2 protein was used to assessed by a commonly used routine diagnostic techniques. It is often called immunohistochemistry. If they did not fully confirm HER-2 protein identification, they again confirmed them by another commonly used technique FISH/SISH.

At first, blood samples from all included patients were collected at the same time of the day after an overnight fasting. However, same procedures were used to get clinical, demographic and anthropometric data from all the included patients by the same research physician and during the first consultation. Patients’ age, weight, height, and menopausal status (i.e., this status expressed whether she was at least 12 months after menopause or reported a bilateral oophorectomy) were collected in this stage.

### Machine learning

The primary objective of this study was to select potential prognostic factors to predict BC. However, the entire prediction model was divided into four steps-

First, data preprocessing that is consist of data cleaning, missing data handling, data integration, data transformation, data reduction, and discretization. Second, to identify the most potential variables which are important for effect and reliable prediction model. Third, the classification models building. Finally, the classification model evaluation using ROC, sensitivity, specificity, positive predictive value, negative predictive value. Figure 1 shows the entire process of our prediction model.

**Figure 1:**
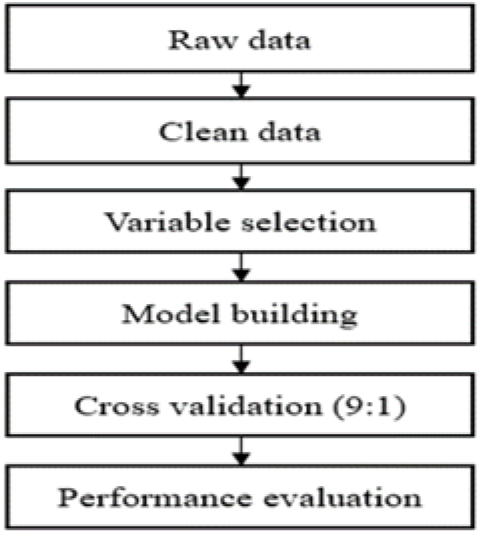
Schematic diagram of machine learning model.

#### Data preprocessing

All those variables were removed that contained >50% missing value. If the variable contained < 5%; we then filled it up with mean value. It a very common and widely used approach dealing missing data in machine learning.

#### Important variables selection

In the real-dataset, some variables are nothing but noise. We selected variables that make sense and actually helps in predicting breast cancer. We used the Boruta package which is a feature ranking and selection algorithm based on random forests algorithm. The advantage of this is that it clearly decides if a variable is important or not and helps to select variables that are statistically significant. **Figure 2** shows the potential variables in the prediction model.

**Figure 2:**
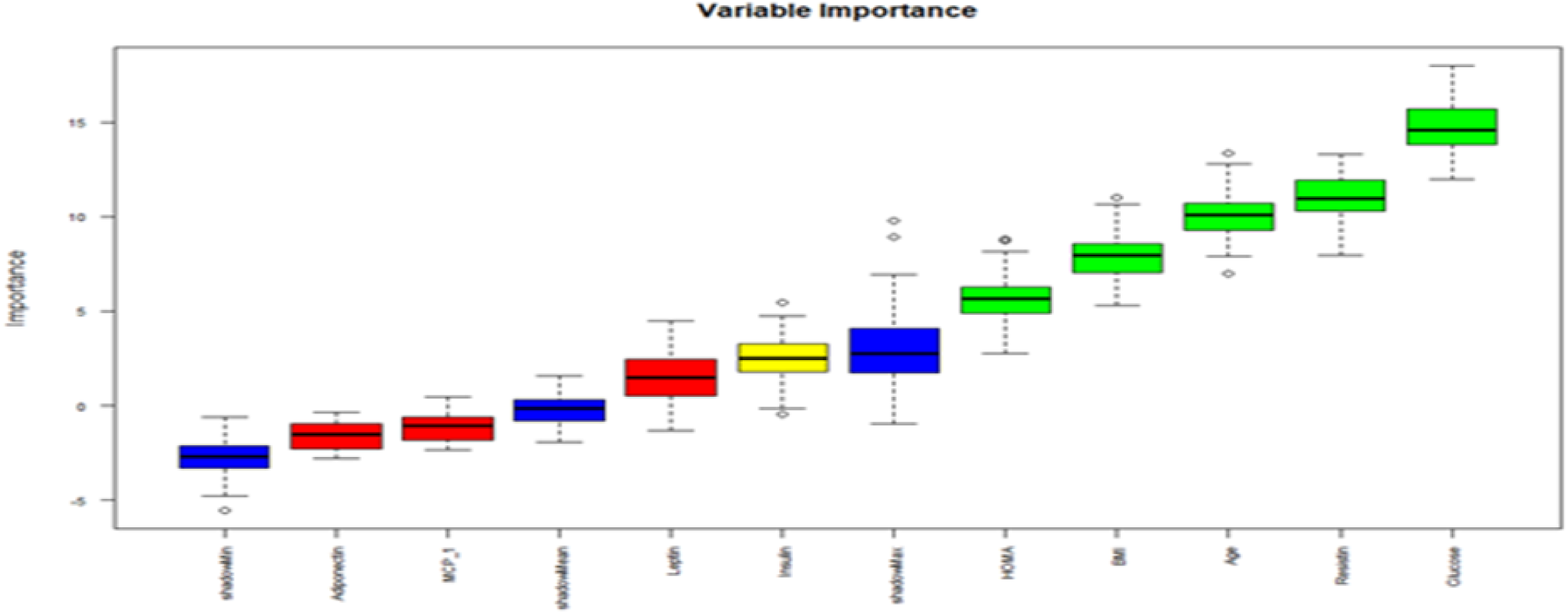
Ranking of potential variables.

**Figure 3:**
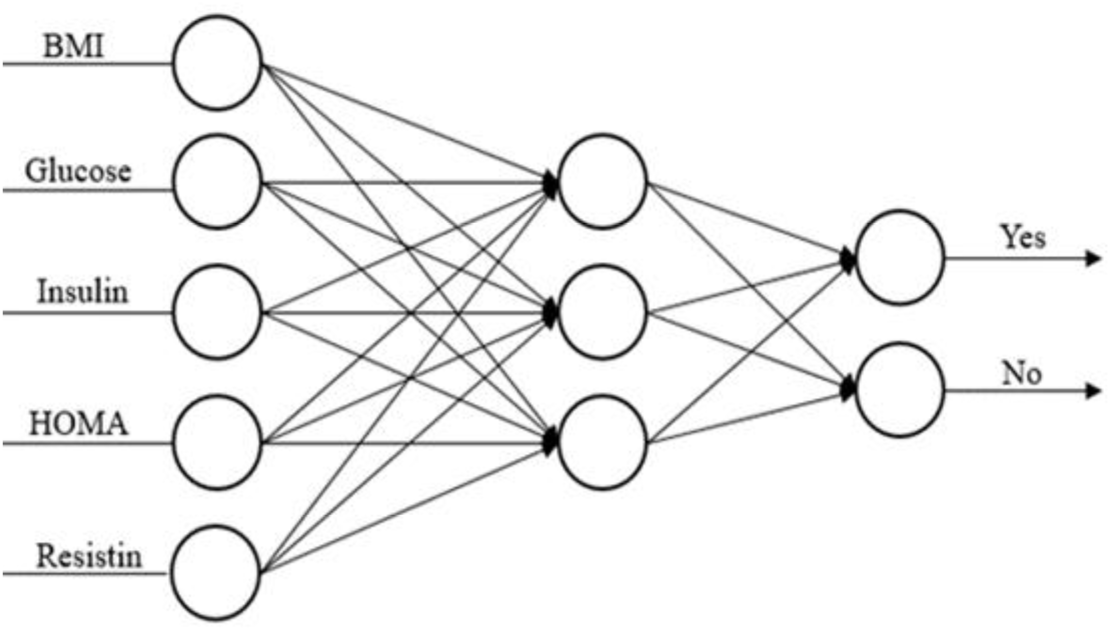
Architecture of ANN for predicting breast cancer.

#### Model building

Six classification models such as Random forest (RF), artificial neural network (ANN), Decision tree (DT), support vector machine (SVM), and logistic regression (LR), K-nearest neighbors (KNN) were used to predict BC patients. These models are very simple, fast and easy to explain. Indeed, these models perform very well when the number of the data sample is not quite big and are robust to overfitting.

An analysis was undertaken in several meaningful stages. We used all important variables in the decision tree, random forest, and support vector machine model. The training set was used to screen out the predictors using univariate logistic regression and developed an ANN prediction model. Stepwise logistic regression was performed to identify the significant predictors of BC. It searches for the best possible regression model by iteratively selecting and dropping variables to arrive at a model with the lowest possible AIC. In this study, it was not so complicated (< 20 vars.), and did just a simple stepwise in ‘both’ directions. Finally, from a pool of shortlisted features, run a full stepwise model to get the final set of selected features

**Table 1** shows the five significant predictors of BC selected by univariate logistic regression. The following five factors are significantly associated with the increased risk of BC: BMI (OR=1.16), HOMA (OR=1.80). The occurrence of BC is inversely related to three predictors, including glucose (OR=0.91), insulin (OR=0.81), and resistin (OR=0.93). Rest of the variables were not significantly associated with BC.

**Table 1:** Predictors of BC analyzed by univariate logistic regression.

| Variable | B | SE | P | OR (95% CI) |
| --- | --- | --- | --- | --- |
| BMI | -2.65 | 1.18 | 0.01 | 1.16(1.06-1.29) |
| Glucose | 1.37 | 3.10 | <0.001 | 0.91(0.85-9.70) |
| Insulin | 4.49 | 1.43 | 0.001 | 0.81(0.59-1.48) |
| HOMA | -1.35 | 4.77 | 0.001 | 1.80(0.13-5.42) |
| Resistin | 7.73 | 3.82 | 0.01 | 0.93(0.87-9.82) |

#### Cross-validation

The model validation technique, 10-fold cross-validation was used assessed models performance and estimation of general error during the whole machine process. In this process, entire dataset divides into equal 10 folds which are approximately the same number of events. Nine-folds use as the training set, and the remaining 1 fold as the testing set. It continues until each fold use once for validation. The results from 10 times validation models are then combined to provide a measure of the overall performance. This process was again repeated 5 times for reducing error.

### Statistical analysis

Descriptive characteristics of the study population were provided including continuous variables as a mean ± standard deviation. The receiver-operating curve was used to assess the performance of these model. R software (version 3.4.2) was used to analyze the basic statistical problem and to construct the ANN prediction model. R is well known as a very powerful prediction tool which contains a collection of visualization tools. However, it also contains the graphical user interface for easily performing algorithms. All statistical tests were two-tailed and p<0.05 was considered significant.

### Model evaluation

The confuse matrix has widely been used for summarizing the performance of the classification model.

#### Accuracy

Accuracy of a model is defined as the total positive instances of the model are divided by the total number of instances. Accuracy parameter provides the percentage of correctly classified instances. The accuracy of model is defined as

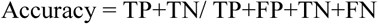

#### Sensitivity

This parameter is used to determine the degree of the attribute to correctly classify the person with diseases and is defined as

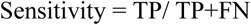

#### Specificity

This parameter is used to determine the degree of the attribute to correctly classify the person without diseases and is defined as

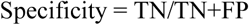

The sensitivity, specificity is also known as quality parameters and used to define the quality of the predicted class. To determine the goodness of the medical diagnosis model basically three parameters are used, these three parameters are accuracy, sensitivity and specificity.

## Results

### Study cohort

Initially, 154 subjects were included in this study. However, 38 subjects were excluded because included patients had BMI above 40 kg/m2 and patients had not at least one of the quantitative variables. Finally, a total of 116 patients (64 cases and 52 healthy controls) were included in our study. Of these 64 breast cancer patients, 13 (21.3%) were tumor grade I, 39 (63.9%) were tumor grade II, and 9 (14.8%) were tumor grade III Additionally, in the case of tumor stage and size, 5 patients had stage 0, 29 (45.3%) patients had stage I, 30 (46.9%) patients had stage II, and more than two-thirds of patients, 54 (84.4%) were having tumor size approximately ≤2 cm. Furthermore, 53 (82.8%) patients had oesterogen (ER) positive, and 5 (7.8%) patients had oesterogen negative. Also, 52 (81.3%) patients showed progesterone receptor positive (PR+), and 6 (9.4%) patients showed progesterone negative (PR-).

### Patients characteristics

The quantitative features of patients and healthy controls are described in table 2. The mean age of BC and healthy controls were 56.67±13.49, and 58.07±18.95 years. The mean BMI (26.98±4.62 vs 28.31±5.42), leptin (26.89±19.21 vs. 26.63±19.33), adiponectin (9.88±6.18 vs. 10.32±7.63) was almost similar in both groups. However, the mean values were different in glucose, insulin, HOMA, resistin, and MCP.

**Table 2:**
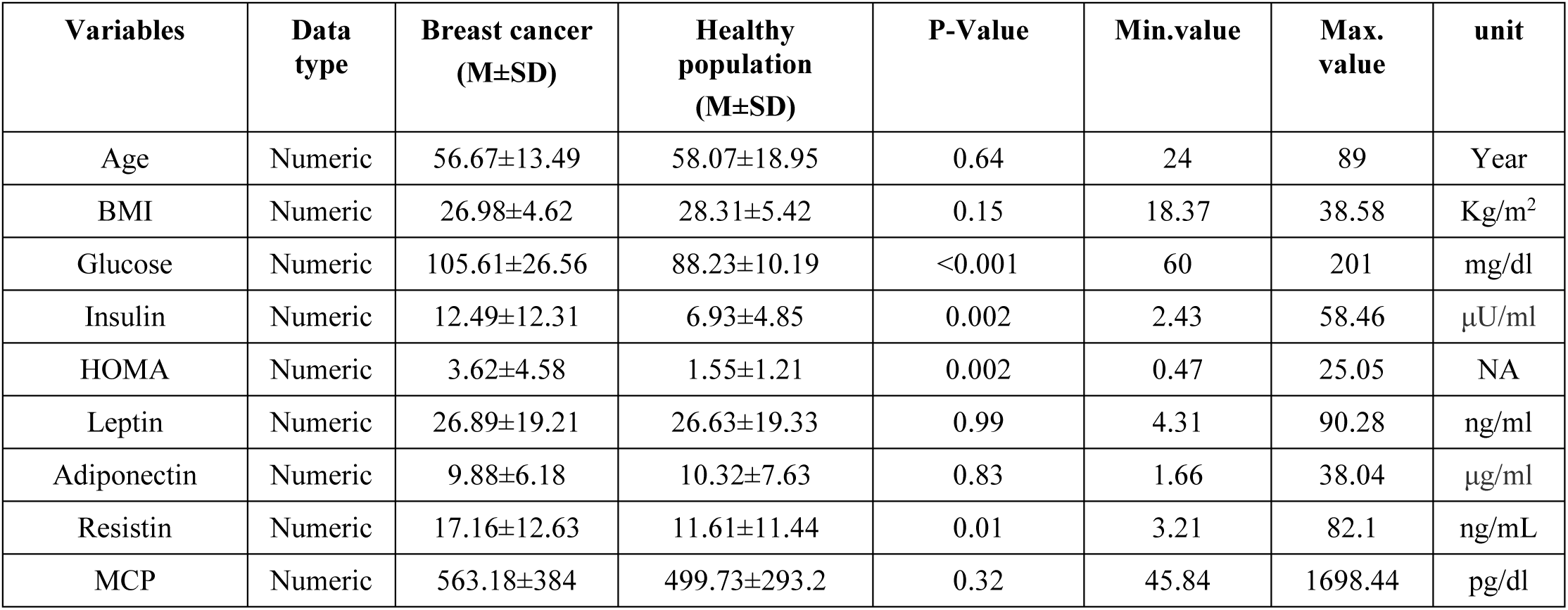
Baseline characteristics of patients.

### Performance evaluation

In **Table 3** the accuracy, sensitivity, specificity, positive predictive value, negative predictive value, and the area under receiving operating curve of six machine learning algorithms are reported. However, KNN and the logistic regression model are superior to DT, RF, SVM, and ANN. Furthermore, KNN exhibited better performance than LR (accuracy: 0.86 vs 0.81; sensitivity: .80 vs 0.70; specificity: 0.91 vs 0.91; PPV: 0.88 vs 0.87; NPV: 0.84 vs 0.78; and AUC: 0.95 vs 0.95). Therefore, it is seemed KNN model to be most suitable for the breast cancer patient’s prediction.

**Table 3:** Performance comparison of different classification models-.

| Models | AUROC | Accuracy | Sensitivity | Specificity | Positive predictive value (PPV) | Negative predictive value (NPV) |
| --- | --- | --- | --- | --- | --- | --- |
| *Decision tree* | 0.71 | 0.72 | 0.70 | 0.75 | 0.70 | 0.75 |
| *Random forest* | 0.80 | 0.72 | 0.83 | 0.60 | 0.71 | 0.75 |
| *Support vector machine* | 0.83 | 0.77 | 0.60 | 0.91 | 0.85 | 0.73 |
| *Logistic regression* | 0.95 | 0.81 | 0.70 | 0.91 | 0.87 | 0.78 |
| *K-nearest neighbors* | 0.95 | 0.86 | 0.80 | 0.91 | 0.88 | 0.84 |
| *Artificial neural network* | 0.92 | 0.80 | 0.83 | 0.72 | 0.76 | 0.80 |

## Discussion

### Main findings

Breast cancer is a major public health concern worldwide. A barrier to timely recognition and management of breast cancer is the long clinically silent phase of the disease. Early diagnosis and timely screening would help to increase the quality of a patient’s care. Therefore, there is a great need for the ability to predict the treatment outcome to be used in the proper and timely treatment decision. However, there is no exact tool in the clinical setting that is designed to distinguish breast cancer and non-breast cancer patients with higher accuracy. In this present study, we tried to developed and assessed prediction models which are based on the machine learning algorithms and data were collected from routine blood analysis. However, this variable could be used as a biomarker of breast cancer. KNN showed superior to the other algorithms used in this study. These results provide the evidence that a machine learning algorithms can discriminate between a patient with breast cancer or without using breast cancer using only 9 variables.

### Interpretation of predictors

The feature selection process that identified most potential variables are clinically sound. These risk factors of BC reported in previous epidemiological studies. However, traditional risk factors of breast cancer remained highly important. Glucose, age, BMI, resistin, HOMA were 5 predictors of top importance. Actually, these predictors make sense like age, BMI, HOMA are considered common risk factors for breast cancer [5] while glucose, resistin is correlated with breast cancer [8, 9]. Additionally, Santillan-Benitez JG, et al. [6] reported in 2013; they used BMI, leptin, CA15-3 and the ratio between Leptin and Adiponectin together to assess as a biomarker for breast cancer and achieved model specificity (80%) and the sensitivity (83.3%). Also, Dalamaga et al. [10] evaluated serum Resistin as a predictor of postmenopausal breast cancer and found an AUC value of 0.72, 95% CI [0.64, 0.79].

### Clinical implication

Breast cancer is the most common type of cancer in women worldwide and the leading cause of cancer death in women. However, it is highly heterogeneous that can drive various clinical outcomes including prognosis, response to treatment and chances of recurrence and metastasis [11]. An important task in personalized medicine is to find out proper biomarkers/risk factors in order to provide the most effective treatment. Early prediction of breast cancer would help to improve survival and quality of life since the disease is generally not curable. We developed and assessed a prediction model and achieved higher accuracy, sensitivity, and specificity. One nine variables were used which potentially be used as a biomarker of breast cancer and can be gathered in routine blood analysis. Therefore, our prediction model would help healthcare providers and organization at multiple levels. Physicians can be used as an assistance tool for proper decision making, and healthcare manager could be stratified patients by using risk factors that would help for budget planning, as high-risk patients tend to require more resource.

### Limitations

This study has several limitations. First, only nine variables were evaluated to predict breast cancer, and could not provide prediction according to the stage of cancer. Second, only one hospital data was used in this study that might be considered as a hindrance of generalization of this prediction model. Third, we did not provide any measurement according to the strength of the variable. Finally, the small sample size was included in this study that might cause over-fitting and may lead to artificially higher accuracy results; although a repeated cross-validation was used to minimize bias; it could not give a guarantee to fully eliminate it.

## Conclusion

The findings of this study suggest that machine learning classification model has potential to predict breast cancer risk from clinical and laboratory variables. However, current classification model needs to be further improved before using real clinical setting. Although, the higher sensitivity, specificity, positive predictive value and negative predictive value of KNN model might have potential to provide useful information for the clinical practice when stratifying breast cancer patients.

## Conflict of interest

None

## References

1. Fitzmaurice C, Dicker D, Pain A, Hamavid H, Moradi-Lakeh M, MacIntyre MF, Allen C, Hansen G, Woodbrook R, Wolfe C (2015) The global burden of cancer 2013. JAMA oncology 1:505–527

2. Corbex M, Burton R, Sancho-Garnier H (2012) Breast cancer early detection methods for low and middle income countries, a review of the evidence. The Breast 21:428–434

3. Slamon DJ, Leyland-Jones B, Shak S, Fuchs H, Paton V, Bajamonde A, Fleming T, Eiermann W, Wolter J, Pegram M (2001) Use of chemotherapy plus a monoclonal antibody against HER2 for metastatic breast cancer that overexpresses HER2. New England Journal of Medicine 344:783–792

4. Smith RA, Saslow D, Sawyer KA, Burke W, Costanza ME, Evans WP, Foster RS, Hendrick E, Eyre HJ, Sener S (2003) American Cancer Society guidelines for breast cancer screening: update 2003. CA: a cancer journal for clinicians 53:141–169

5. Crisóstomo J, Matafome P, Santos-Silva D, Gomes AL, Gomes M, Patrício M, Letra L, Sarmento-Ribeiro AB, Santos L, Seiça R (2016) Hyperresistinemia and metabolic dysregulation: a risky crosstalk in obese breast cancer. Endocrine 53:433–442

6. Santillán-Benítez JG, Mendieta-Zerón H, Gómez-Oliván LM, Torres-Juárez JJ, González-Bañales JM, Hernández-Peña LV, Ordóñez-Quiroz A (2013) The Tetrad BMI, Leptin, Leptin/Adiponectin (L/A) Ratio and CA 15-3 are Reliable Biomarkers of Breast Cancer. Journal of clinical laboratory analysis 27:12–20

7. Patrício M, Pereira J, Crisóstomo J, Matafome P, Gomes M, Seiça R, Caramelo F (2018) Using Resistin, glucose, age and BMI to predict the presence of breast cancer. BMC cancer 18:29

8. Kang J-H, Yu B-Y, Youn D-S (2007) Relationship of serum adiponectin and resistin levels with breast cancer risk. Journal of Korean medical science 22:117–121

9. Muti P, Quattrin T, Grant BJ, Krogh V, Micheli A, Schünemann HJ, Ram M, Freudenheim JL, Sieri S, Trevisan M (2002) Fasting glucose is a risk factor for breast cancer: a prospective study. Cancer Epidemiology and Prevention Biomarkers 11:1361–1368

10. Dalamaga M, Sotiropoulos G, Karmaniolas K, Pelekanos N, Papadavid E, Lekka A (2013) Serum resistin: a biomarker of breast cancer in postmenopausal women? Association with clinicopathological characteristics, tumor markers, inflammatory and metabolic parameters. Clinical biochemistry 46:584–590

11. Wang C, Machiraju R, Huang K (2014) Breast cancer patient stratification using a molecular regularized consensus clustering method. Methods 67:304–312

